# Spatiotemporal dynamics and selectivity of mRNA translation during mouse pre-implantation development

**DOI:** 10.1101/2024.10.28.620693

**Authors:** Hao Ming, Rajan Iyyappan, Kianoush Kakavand, Michal Dvoran, Andrej Susor, Zongliang Jiang

## Abstract

Translational regulation is pivotal during preimplantation development. However, how mRNAs are selected for temporal regulation and their dynamic utilization and fate during this period are still unknown. Using a high-resolution ribosome profiling approach, we analyzed the transcriptome, as well as monosome- and polysome-bound RNAs of mouse oocytes and embryos, defining an unprecedented extent of spatiotemporal translational landscapes during this rapid developmental phase. We observed previously unknown mechanisms of translational selectivity, i.e., stage-wise deferral of loading monosome-bound mRNAs to polysome for active translation, continuous translation of both monosome and polysome-bound mRNAs at the same developmental stage, and priming to monosomes after initial activation. We showed that a eukaryotic initiation factor Eif1ad3, which is exclusively translated in the 2-Cell embryo, is required for ribosome biogenesis post embryonic genome activation. Our study thus provides genome-wide datasets and analyses of spatiotemporal translational dynamics accompanying mammalian germ cell and embryonic development and reveals the contribution of a novel translation initiation factor to mammalian pre-implantation development.

## INTRODUCTION

Translational control of mRNA is vital across various cellular processes, enabling rapid protein synthesis essential for both immediate cellular needs and longer-term physiological changes ^1,2^. During mammalian pre-implantation development, the degradation of maternal stored mRNAs and activation of embryonic genome are precisely regulated, in large part by the translational and posttranslational regulation ^3^. For example, fully grown oocytes rely entirely on the translation of stored mRNA for maturation and fertilization, given their lack of active transcription ^4^. However, a central gap in our understanding of post-transcriptional regulation exists, particularly how mRNAs are selected for spatial and temporal regulation and their dynamics of mRNA utilization and fate during oocyte maturation, fertilization, embryonic genome activation (EGA) and early differentiation. Thus, understanding how mRNA is selectively translated during these critical transitions is essential for elucidating the mechanisms underlying successful embryo development.

In the past decade, the dynamics of genome-wide transcription, translation, and protein expression during mammalian pre-implantation embryo development have been characterized, however, limitations remain. First, the mRNA expression from transcriptomic profiling do not represent their functional status ^5^. Second, despite of proteomic analysis of oocytes and embryos has been achieved in several mammalian species by mass spectrometry ^6–9^, the methodology faces challenges such as low sensitivity and limited coverage due to the scarcity of material available. Third, recent studies have developed low-input ribosome profiling approach (LiRibo-seq and Ribo-ITP) and provided the first translational dynamics of mouse pre-implantation development ^10,11^. However, these studies were confined to an analysis of ribosome bound mRNAs as a whole, while ignoring the variation encountered in the different numbers of ribosome-bound mRNAs and how the specific mRNAs are preferentially selected for translation, underscoring the need for improved techniques to better understand the dynamics of mRNA usage and fate during these crucial stages.

To decipher comprehensive translational regulation in mouse pre-implantation development, we have employed an optimized high-resolution single-fraction of Scarce Sample Polysome Profiling (SSP-profiling) ^12^ on mouse oocytes and pre-implantation embryos. This methodology has successfully provided systematic monitoring of non-translated RNA (free mRNA), mRNA prepared for translation (different numbers of monosomes-bound mRNA), and actively translating mRNA (different numbers of polysomes-bound mRNA) in a bovine embryo model^13^. Therefore, here we have provided a high-resolution and genome-wide spatial translational dynamics during mouse pre-implantation embryo development. This resource allows to dissect the dynamic of specific mRNA that is untranslated, translated, or degraded and the translation selectivity of individual mRNA during early embryogenesis, therefore adding the missing picture of translational regulation of pre-implantation development. More importantly, our findings highlight the role of a translation initiation factor eIF1ad3 and its variants that are translated exclusively at the 2-Cell embryos. This stage-specific expression suggests a vital role of these factors in regulating protein synthesis immediately following major genome activation. The presence of eIF1ad3 at this juncture indicates its importance in the developmental transitions that characterize continuum of early embryogenesis, providing a clearer understanding of the molecular underpinnings that drive crucial phases of development.

## RESULTS

### High-resolution ribosome profiling of mouse oocytes and pre-implantation embryos

By employing an high-resolution ribosome profiling approach, an improved SSP-profiling protocol followed by RNA-seq as described previously ^12,13^, we analyzed translational profiles of mRNAs that are bounded on different number of translating ribosomes in mouse oocytes at the germinal vesicle (GV) and metaphase II (MII) stages, as well as pre-implantation embryos at the zygote, 2-Cell, 4-Cell, 8-Cell, morula, and blastocyst stages (**Fig. 1A**). For each stage, 200 oocytes or embryos were utilized, and the experiment was performed with three biological replicates per stage. Following ultracentrifugation on sucrose gradients, ten equal volumes of fractions (F1-F10) from each developmental stage were collected for low-input RNA-seq analysis (methods), providing a high-resolution mRNA translational profiles of pre-implantation development (**Fig. 1A**). To systematically explore the correlation of global mRNA transcription and translation, RNA-seq analysis of total mRNA was conducted with 20 oocytes (GV and MII stages) (n= 4) and 20 embryos (n=4) at each developmental stages collected from the same batches used for high-resolution ribosome profiling (**Fig. 1A**). Reproducibility of sequencing datasets was confirmed by principal component analysis (PCA), showing that all replicate samples from the same biological groups consistently aligned with each other (**Fig. 1B**, **Fig. S1A, B**). The ten fractions (F1-F10) were categorized into three groups: fractions F1-F2 primarily containing free RNA, F3-F5 predominantly binding monosome, and F6-F10 mainly binding polysome (**Fig. S1B**), which is consistent with what we’ve described in pre-implantation development of bovine embryos ^13^.

**Figure 1.**
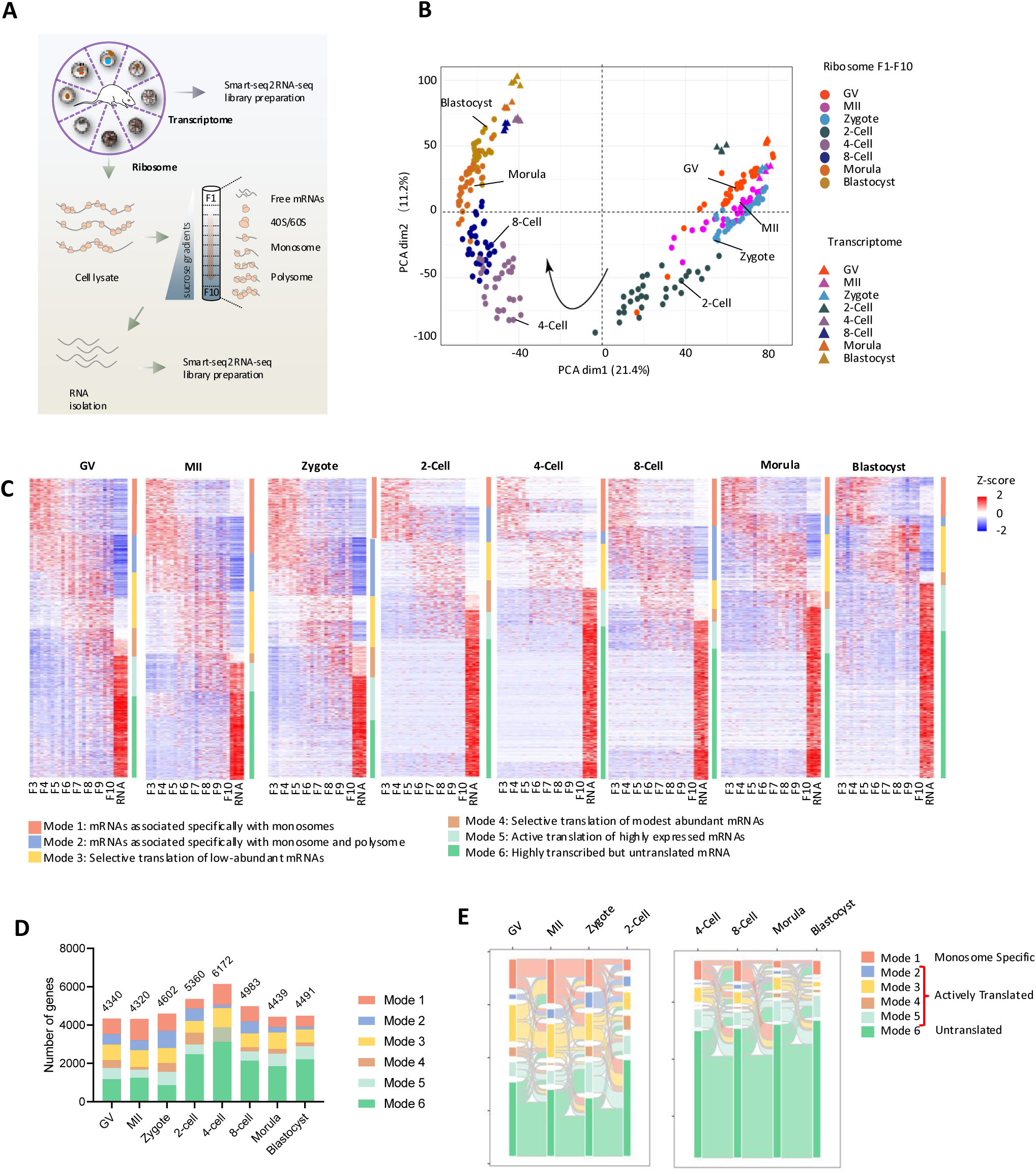
High-resolution ribosome profiling of mouse oocytes and pre-implantation embryos. **A**. Scheme of high-resolution ribosome profiling of mouse pre-implantation development. **B**. Principal component analysis (PCA) of translatome (F1-F10) (n=3) and transcriptomes (n=4) of mouse oocytes and pre-implantation embryos. **C**. Heatmap showing six diverse modes of translational selectivity during mouse oocyte and pre-implantation development. The color spectrum, ranging from red through white to blue, indicates high to low levels of gene expression. Mode 1: mRNAs associated specifically with monosomes (red bar); Mode 2: mRNAs associated specifically with monosome and polysome (blue bar); Mode 3: Selective translation of low-abundant mRNAs (yellow bar); Mode 4: Selective translation of modest abundant mRNAs (brown bar); Mode 5: Active translation of highly expressed mRNAs (cyan bar); Mode 6: Highly transcribed but untranslated mRNA (green bar). **D**. Number of genes from different modes across all developmental stages. **E.** Sankey diagram showing the flow of genes in different modes across developmental stages.

We found that the GV and MII oocytes, and zygotes were clustered together, while a distinct separation between them and 2-Cell stage, when major embryonic genome activation (EGA) occurs ^14,15^ (**Fig. 1B**, **Fig. S1A**). Subsequently, a marked shift was observed in 4-Cell embryos and the stages from 4-Cell onwards showed a continuous but steady progression up to the blastocyst stage (**Fig. 1B, Fig.**).**S** T**1**h**A**e PCA plot also indicated distinct separation between polysome-bound mRNA and monosome-bound mRNA at each stage, highlighting that despite of both being ribosome-bound, a clear gap exists. Interestingly, a significant variance between the different ribosomal fractions was observed at the 2-Cell stage, which decreased at later stages (**Fig. S1A**), possibly reflecting the selective gene translation associated with EGA. In addition, a clear difference between transcriptome and translatome profiles was evident across all developmental stages (**Fig. 1B**), indicating that the transcriptome data do not reflect real-time mRNA translation, as has been found in other studies ^13,16,17^.

### Diverse modes of translational selectivity during mouse pre-implantation development

To analyze the complex interplay between transcriptional and translational changes during mouse pre-implantation embryo development, we employed the *fuzzy k-means* clustering algorithm on datasets of whole transcriptomes, monosome-occupied, and polysome-occupied mRNAs. We identified six modes of translational selectivity (mRNA association with ribosomes) in each developmental stage (**Fig. 1C, Table. S1**): Mode 1: mRNAs associated specifically with monosomes; Mode 2: mRNAs associated with both monosome and polysome; Mode 3: Selective translation of low-abundant mRNAs; Mode 4: Selective translation of modest abundant mRNAs; Mode 5: Active translation of highly expressed mRNAs; Mode 6: Highly transcribed but untranslated mRNAs.

Despite the differences in mRNA occupancy at all developmental stages, the total number of genes we identified at each stage did not change significantly except at the 2- and 4-Cell stages, indicating a boost in transcription during EGA (**Fig. 1D**). Given the marked shift of transcriptional and translational landscapes during EGA at the 2-Cell stage, we explored the flow of genes before and after EGA separately and found that different modes of categorized mRNAs were occupied differently during oocyte maturation and early embryo development (**Fig. 1E**). To simplify the analysis, we categorized mRNAs into three broad groups based on their ribosomal occupation to better understand their roles in oocyte-zygotic transition and embryogenesis: 1) Monosome specific (Mode1), 2) Actively translated (Modes 2, 3, 4, 5) and 3) Untranslated (Mode 6) **(Fig. 1E)**.

The actively translated group showed a stage-specific progression of gene ontology (GO) terms, reflecting shifts in biological processes necessary for each stage of development (**Table 1**). The fluctuation in the number of genes within these modes also indicated significant changes in biological processes and cellular functions from one developmental stage to the next. Notably, genes initially found in monosome-specific or untranslated modes often transitioned into actively translated modes in subsequent stages, underscoring their importance in developmental progression. For example, *Cdk2* and *Apob* showcased significant shifts in their translational profiles at critical stages, aligning with their roles in cell cycle regulation ^18^ and nutrient supply ^19,20^, respectively (**Fig. S2A, B**). Additionally, a considerable share of mRNA stopped being translated while maintaining a high level in the sequential stage (from mode 3/4/5 to mode 6), such as *Mreg* and *Sesn1* at 2-Cell stage. These genes were presumed to be either useful for later developmental stage or degraded gradually (**Fig. S2C, D**). This dynamic translation landscape illustrates the adaptive translation mechanisms that are essential for the developmental transitions within the embryonic stages and ensure that the physiological and morphological changes are precisely tailored to the needs of the embryo at each stage.

**Table 1.**
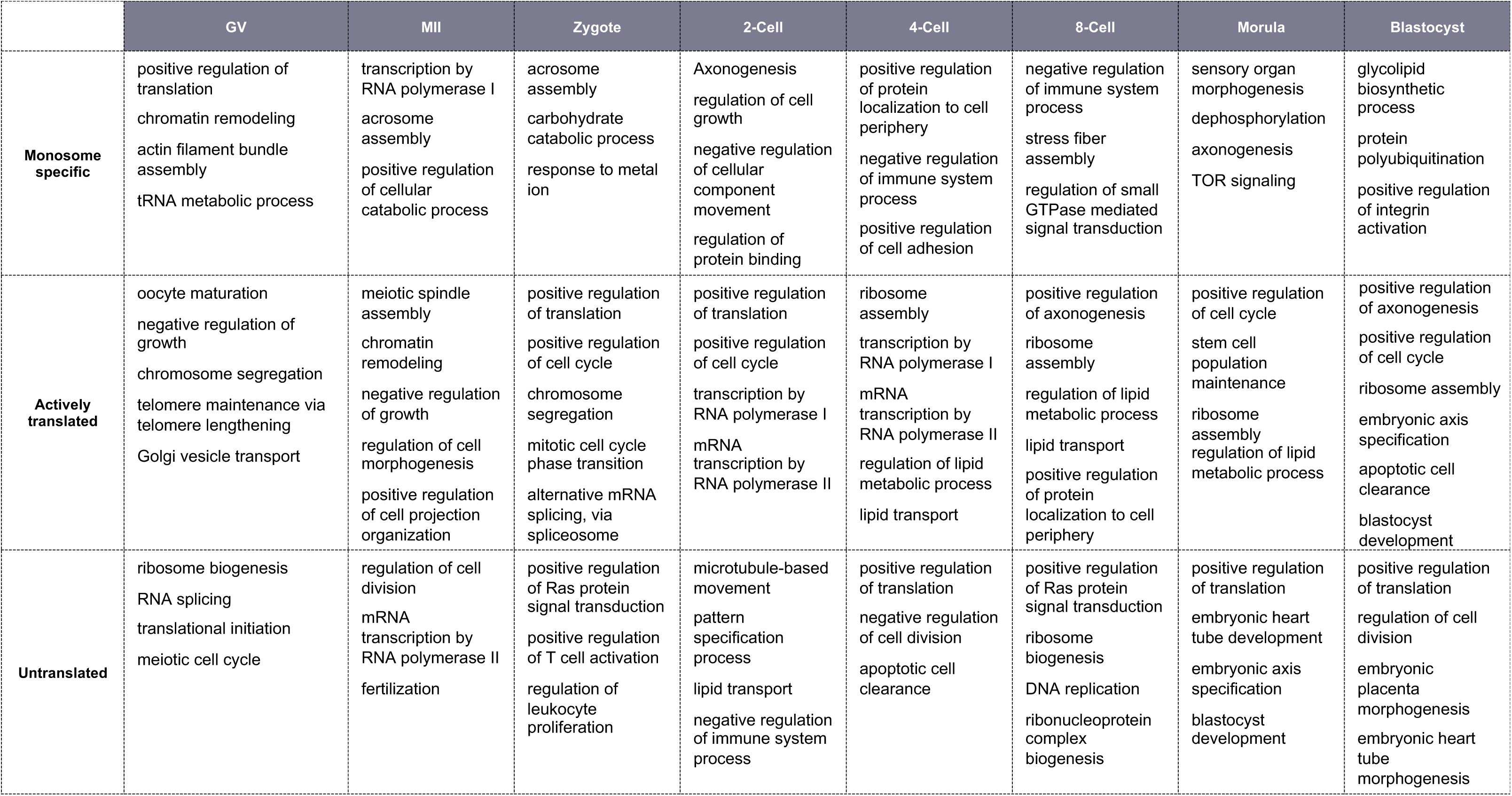
Top enriched GO terms associated with the genes from identified six modes of translational selectivity in each developmental stage of mouse pre-implantation development.

### The dynamics of mRNA translational fate during mouse pre-implantation development

To explore the dynamics of specific mRNA that bounded to different number of ribosomes (termed mRNA translational fate) across different developmental stages, we refined our analysis by applying the *fuzzy k-means algorithm* to categorize profiles of free RNA (fractions F1-F2), monosome-occupied RNA (Fractions F3-F5), and polysome-occupied RNA (Fractions F6-F10) separately (**Fig. 2A**). This detailed clustering was conducted for each stage and is presented in **Fig. 2B-D**. This method enabled us to characterize the changes of RNA usage throughout development, offering a clearer insight into the activation and suppression of genes as development advances. Based on the expression and usage patterns, we clustered them into twelve distinct groups (**Fig. 2B-D**). We also confirmed that oocyte marker genes such as *Zar1*, *Ooep*, and *Mos* were specifically enriched at the oocyte stage while trophectoderm markers such as *Eomes*, *Krt8*, and *Cdx2* were specific enriched at the blastocyst stage, demonstrating the consistent trends across datasets (**Fig. S3A**).

**Figure 2.**
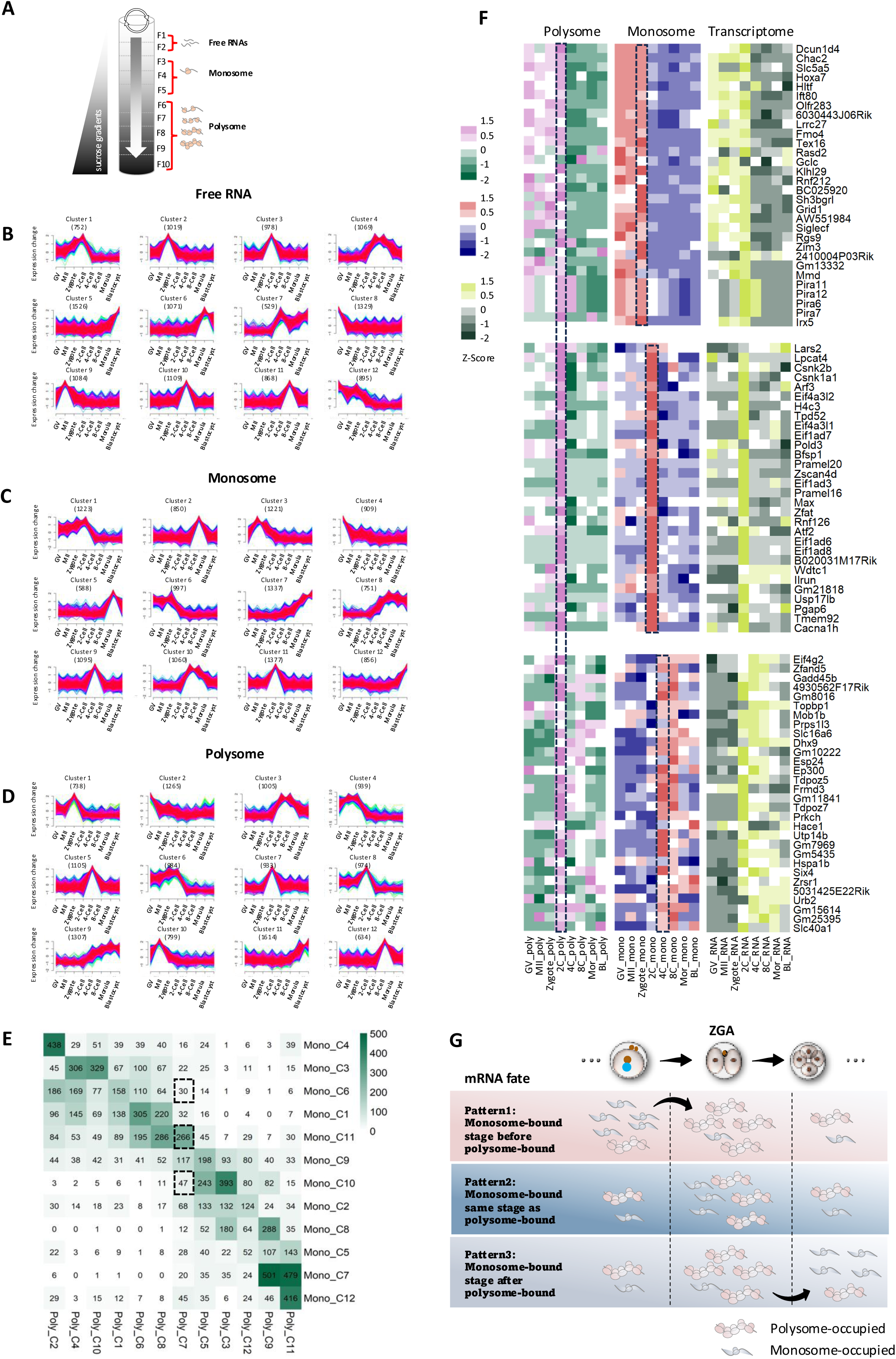
The dynamics of mRNA translational fate during mouse pre-implantation development. **A**. Scheme of the fractions from top to bottom assembled as free RNA (F1-F2), monosome-occupied RNA (F3-F5), and polysome-occupied RNA (F6-F10). **B-D.** Fuzzy c-means clustering identified twelve distinct temporal patterns of free RNA (**B**), monosome-bound mRNA (**C**), and polysome-bound mRNA. (**D**). The ‘x’ axis represents developmental stages in time course, while the ‘y’ axis represents log2-transformed, normalized intensity ratios in each stage. **E.** The heatmap showing the number of genes overlapped from the identified stage-specific clusters between monosome- and polysome layers based on Figure 2C**, D**. The color scheme, from white to green, indicates the number of intersection genes from low to high. **F.** Heatmap showing the expression levels of specific genes across the polysome, monosome, and transcriptome profiles. Top panel: Genes peaking at the 2-Cell stage in the polysome profile, while peaking at the zygote stage in the monosome profile. Middle panel: Genes peaking at the 2-Cell stage in the polysome profile, while peaking at the 4-Cell stage in the monosome profile. Bottom panel: Genes peaking at the 2-Cell stage in both the polysome and monosome profiles. **G.** An illustration graph summarizing three different mRNA fates during early embryo development, i.e., mRNAs are monosome-bound stage before polysome-bound (top panel); or monosome-bound same stage as polysome-bound (middle panel); or monosome-bound stage after polysome-bound (bottom panel). Example given here is focusing on two cell stage.

While most genes showed stage-specific expression, many monosome-bound genes either remained as such for later use or transitioned to polysome-bound to support immediate protein synthesis for specific stages. This variation illustrates how maternally stored mRNAs are regulated based on developmental needs (**Fig. 2E**). GO analysis of stage-specific genes revealed the dominant biological functions for each RNA profile (**Fig. S3C, D)**, showing consistent functions across the different RNA profiles. For example, GO terms for microtubule-based movement that are associated with the monosome at the GV and MII oocytes shifted to the zygotes and 2-Cells in the polysome profiles. Similarly, ribosome biogenesis and mRNA splicing, which were prominent in the monosome profiles at the 4- and 8-Cells, were more prominent in the polysome profiles at the morulae, mainly due to the differences in non-overlapping genes between the monosome and polysome profiles (**Fig. S3C, D**).

Given the EGA triggers extensive gene activation and translation, we specifically focused on the mRNA translational fate in the 2-Cell embryos. We found that genes in polysome-bound (Clusters 6, 7 and 8) are actively translated, especially at the 2-Cell stage (**Fig. 2D; Fig. S3B**). We also noticed three distinct patterns: 1) mRNAs are initially not translationally active, becoming highly active during EGA and subsequently returning to a quiescent state (**Fig. 2F top panel**); 2) mRNAs remain low in abundance and are rarely ribosome-bound until a spike in transcription and translation in the 2-Cells (**Fig. 2F middle panel**); and 3) mRNAs are actively translated in the 2-Cells, then transition to monosome-bound status afterwards, either being degraded or reserved for later stages (**Fig. 2F bottom panel**). These patterns demonstrate the linear progression of early embryonic development and emphasize the strategic regulation of mRNA storage and translation, which is critical for the subsequent cellular process in the pre-implantation development (**Fig. 2G**).

We extended our analysis across all stages to investigate the translational fate of mRNAs based on the patterns identified, and found that delayed translational activation was found to occur extensively (**Fig. 2G**). We were able to categorize genes into following three groups: 1) those overlapping between monosome- and polysome-occupied RNA at the same stage, 2) those polysome-occupied one stage after being monosome-occupied, and 3) those polysome-occupied two stages after monosome occupancy. We also explored the biological functions of genes in these groups (**Fig. S4A, B**). For example, the overlapping genes at the same stages regulate essential biological processes sequentially, such as meiosis in oocytes, mitotic cell cycle and transcription in 2-Cells, ribosomal large subunit biogenesis in 4-Cells, and stem cell differentiation in blastocysts. On the other hand, the genes with delayed translational activation suggested their involvement in crucial later embryonic processes. These include mRNA stability regulation in zygotes, mitotic cell cycle phase transition in 2-Cells, RNA and histone modifications in morulae, and Rho protein signal transduction in blastocysts.

To validate this delayed translational activation, we examined several specific genes that are peaked either before or after EGA (**Fig. S5**). For example, *Zbed3*, a key regulator of *Wnt* signaling and necessary for normal zygotic division and organelle distribution ^21–23^, peaked at the MII stage in the free and monosome-occupied layers but continued to increase until the 2-Cell stage in the polysome-occupied group. The RNA methyltransferase NSUN5, known to regulate maternal-to-zygotic transition in mice and plays a key role in cell size and proliferation via its effect on translation ^24,25^, showed a dramatic increase in both monosome- and polysome-bound layers starting at the 2-Cell stage. After the 8-Cell stage, the level in the monosomal layer decreased, while it peaked in polysomal layer at the morula stage. Other validated genes showing delayed activation included *Abca12*, *Brsk2*, and *Jak1* (pre-EGA), and *Rexo4*, *Dgkz*, and *Eps8l2* (post-EGA). We also confirmed delayed translation in free RNAs, such as *Tubb4a*, *Itga7*, *Hdac2*, and *Tdgf1*. Notably, *Tdgf1* (*Cripto-1*) peaked in the free RNA layer at the 4-Cell stage, while peaked in both the monosome and polysome layers at the 8-Cell stage.

Collectively, during pre-implantation development, discrepancies arise not only between the transcriptome and the translatome, but also within the translatome itself, as mRNAs selectively bind to monosomes or polysomes. Oocytes and embryos regulate mRNA translational fate through a dynamic translation process. Some mRNAs are stored on monosomes for later activation, while others can be continuously translated by binding to both monosomes and polysomes at the same stage or by being stored on monosomes after early activation. Our high-resolution polysome profiling enables us to track mRNA translational dynamics and offers deeper insights into translational regulation during early development.

### Translational switch occurs in key developmental transitions

The development and competence of pre-implantation embryos are driven by several key events, including oocyte maturation, EGA, and first lineage segregation from 8-Cell through blastocyst (**Fig. 3A**). These transitions rely heavily on real-time protein synthesis, making precise translation regulation essential. To map the trajectory of translational changes during these critical transitions, we compared both polysome- and monosome-occupied mRNAs at each stage to assess how they are activated or deactivated during oocyte maturation and embryo development (**Fig. 3B**). Particularly, in each of the transitions, we identified groups of genes that are enriched in the monosome before transition, but later bound to polysomes (**Fig. 3A, C, E, G**). GO analysis of these genes suggested that they play crucial roles in these stage-specific transitions (**Fig. 3D, F, H**).

**Figure 3.**
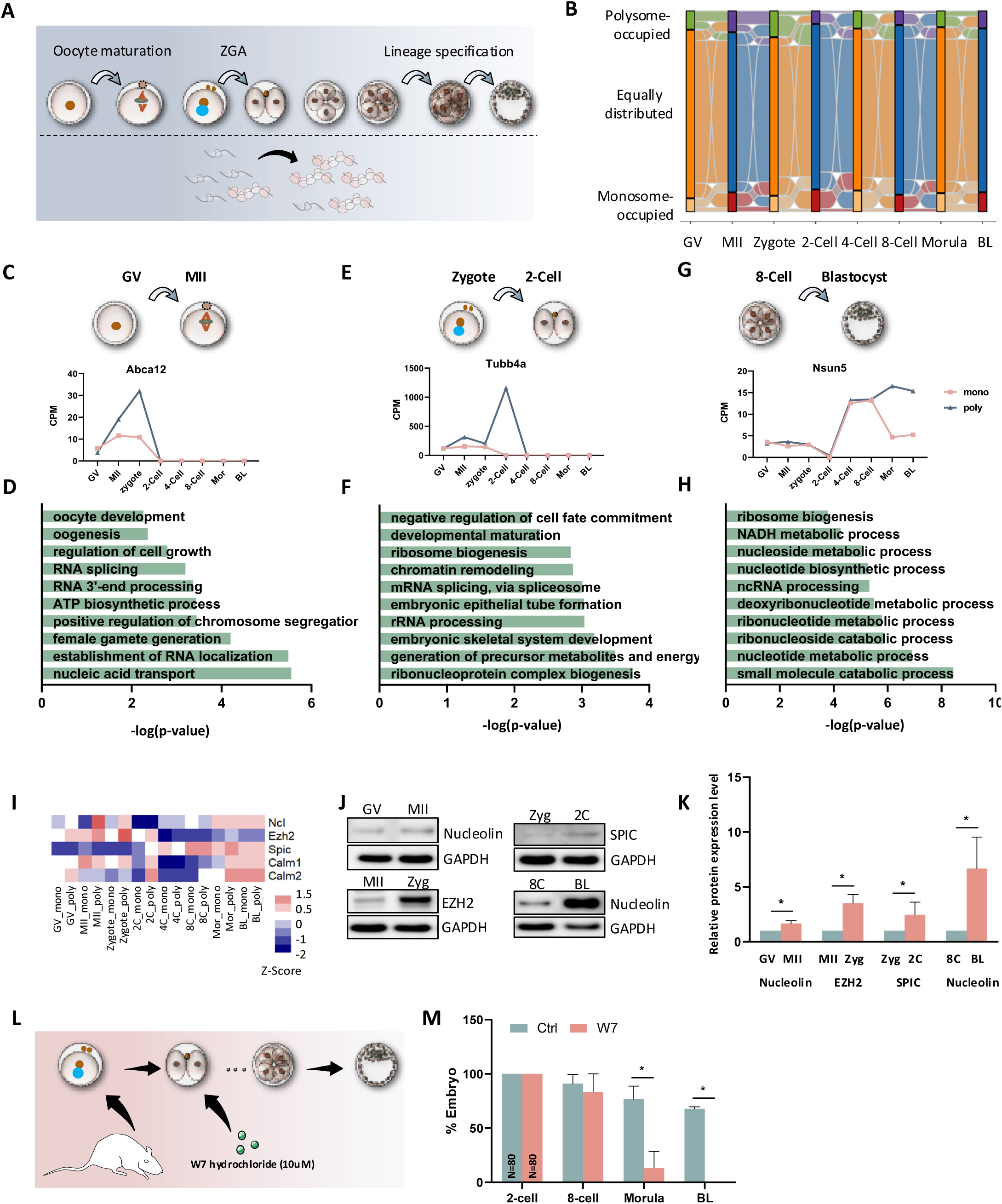
Translational switch occurs in key developmental transitions. **A.** Scheme of the critical developmental transitions during mouse pre-implantation development. **B.** Sankey diagram showing the up- and down-regulated genes (Foldchange > 2, P-value < 0.05) between polysome-occupied mRNAs (F6-F10) and monosome-occupied mRNAs (F3-F5) in each developmental stage. The flow is initiated from GV oocytes, including up-regulated in polysome layers (Top), up-regulated in monosome layers (Bottom), and equally distributed mRNA (not significant bias, Middle). **C-H.** The analysis of genes and their represented top enriched GO terms. Each is enriched in monosome layer in one stage, while enriched in polysome layers in its next developmental stage during GV to MII transition (**C, D**), zygote to 2-Cell transition (**E, F**), and 8-Cell to blastocyst transition (**G, H**). **I**. Heatmap showing the expression level of candidate genes in monosome- and polysome-occupied profiles across different developmental stages. The color spectrum, ranging from red through white to blue, indicates high to low levels of gene expression. Ncl, Nucleolin. **J.** Representative protein levels of selected genes during critical cell fate transitions. GAPDH was used as a loading control. **K.** Quantitative analysis of relative protein levels. **L.** Scheme showing treating 2-Cell embryos with 10mM W7 hydrochloride to detect the effect of Calm-2 inhibition on mouse embryo development. **M.** Quantification of embryo development after inhibition of Calm-2.

To validate these “active genes”, we applied strict cutoffs to prioritize a short list of genes (**Fig. S6**) and employed western blot analysis to validate our sequencing results (**Fig. 3I-K**). To explore stage-specific polysome activation, we focused on *Calm1* and *Calm2*, which encode calmodulin, a key regulator of intracellular Ca^2+^ levels and essential for embryonic development ^26,27^. Both genes peaked at both 2-Cell and morula/blastocyst stages, suggesting their involvement in EGA and lineage specification. It has been reported that calmodulin antagonist W-7 has inhibitory effect on first cleavage during mouse preimplantation development ^28^. To overcome the arrest of first cleavage, we cultured embryos with W-7 after first cleavage and maintained them until the end of culture (**Fig. 3L**). The developmental rate was not affected until morula stage, but none of them could proceed to blastocyst (**Fig. 3M**), highlighting the critical role of *Calm1* and *Calm2* translation in regulating first lineage specification. This underlines significant value of the high-resolution of single-fraction ribosome profiling during critical stages of development.

### Integrated analysis of mRNA active translation with their protein expression, poly(A) tail length, and m6A modifications during mouse pre-implantation development

To explore the correlation between gene translation and its protein expression, we integrated our polysome-associated mRNA profiles with a published proteomics dataset ^7^ at the matched developmental stages. We observed that polysome occupancy and protein expression were not positively correlated at the 4-Cell and 8-Cell stages (**Fig. 4A**). Upon further investigation across stages, we detected a delayed correlation between polysome levels at the 4-/8-Cell stages and protein levels at the morula/blastocyst stages (**Fig. 4B, Fig. S7**), suggesting that during the second and third rounds of cell division, protein maturation is delayed until embryo compaction. Additionally, poly(A) tail length regulates mRNA translation ^29,30^, by integrating datasets of poly(A) tail length ^31^ and polysome occupation from our study, we found a delayed positive correlation between poly(A) tail length at the 2-Cell stage and polysome levels at the 4-/8-Cell stages (**Fig. 4C**). This implies that embryos prepare for translation at later stages through re-adenylation during EGA ^32^. Collectively, these results demonstrate that embryos optimize energy and time by preparing mRNA for rapid translation and functional activation at later stages through re-adenylation or ribosome binding (**Fig. 4D**).

**Figure 4.**
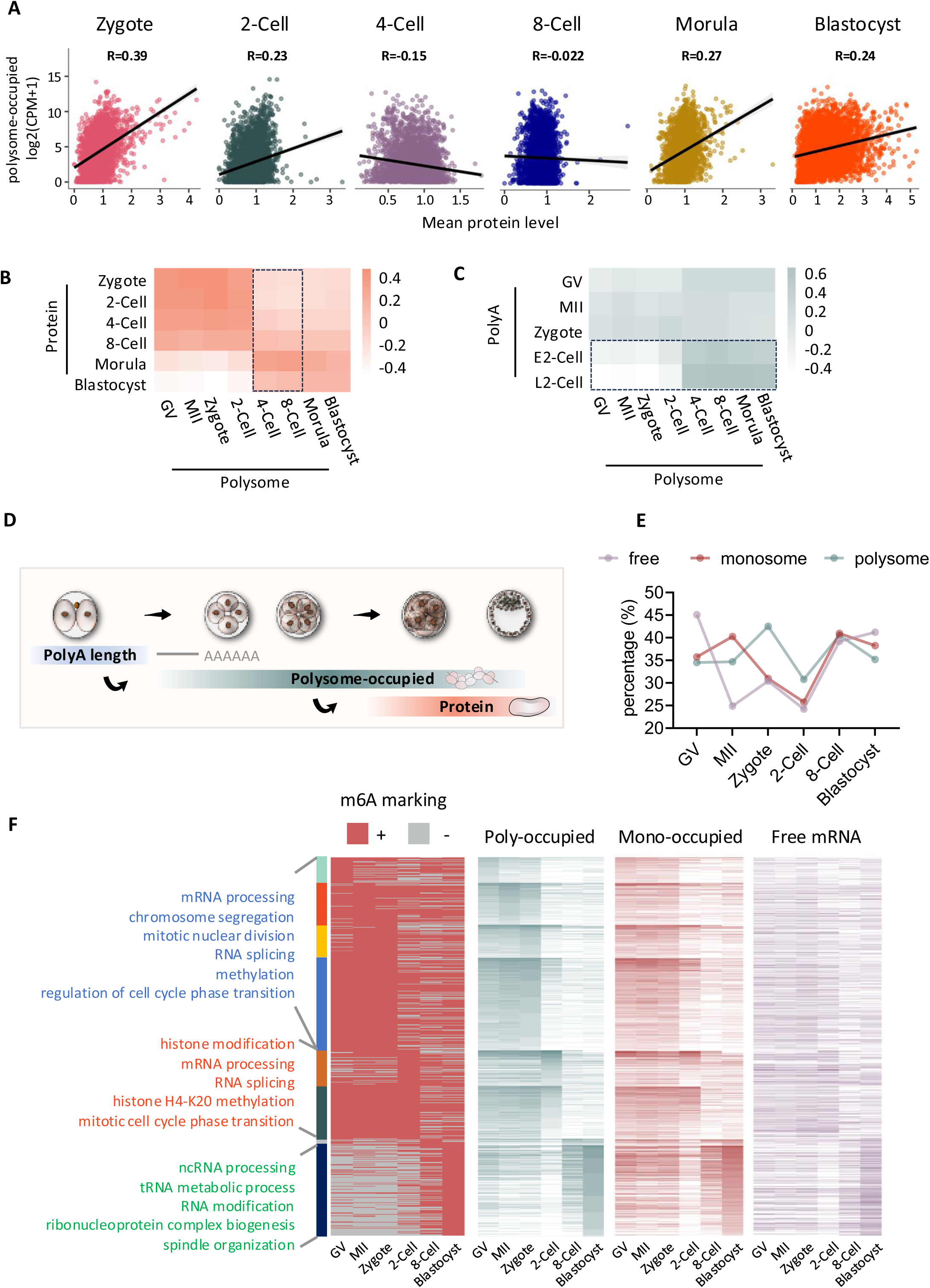
Integrated analysis of mRNA active translation with their protein expression, poly(A) tail length, and m6A modifications during mouse pre-implantation development. **A.** Correlation between polysome-occupied RNA level and protein level at each developmental stage. **B.** Correlation between polysome-occupied RNA level and protein level across developmental stages. **C.** Correlation between polysome-occupied RNA level and poly(A) tail length across developmental stages. **D.** Scheme of the delayed correlation between polysome profile and proteomic, Poly(A) tail length profiles. **E.** Percentage of stage specifically enriched free RNA, monosome-bound, and polysome-bound genes (characterized from Fuzzy c-means clustering results in Figure 2A-C with m6A modification profiles. **F.** Heatmap showing eight groups of genes were identified from integrative analysis of stage-specifically expressed genes from polysome-bound profile with m6A modification profiles. The top GO terms from each group were listed on the left panel. Expression of corresponding genes’ RNA modification profiles and their characterized as poly-occupied, monosome-occupied, and free RNA layers were shown on the right panel.

In addition to these mechanisms, another critical aspect of translation regulation is the role of N6-methyladenosine (m6A), the most prevalent mRNA modification in eukaryotes ^33^. m6A enhances mRNA translation by recruiting ribosomes through the YTHDF1-eIF3 pathway ^34–36^. To investigate whether m6A controls translation during mouse pre-implantation development, we compared the previously published transcriptome-wide m6A modifications dynamics ^37^ with our translational profiles across mouse pre-implantation development. We first examined the percentage of genes at each stage from the free mRNA, monosome-occupied mRNA, and polysome-occupied mRNA profiles that were modified by m6A (**Fig. 4E**). We found that the percentage of m6A modification fluctuates significantly across stages, especially for free and monosome-occupied mRNAs. During the zygote and 2-Cell stages, the percentage of m6A-modified RNA in the polysome layer was higher than in the other two (monosome and free) layers, suggesting a pivotal role for m6A in regulating translation during zygotic gene activation, as previously indicated ^37^. To further understand the relationship between m6A modification and translational activity, we identified genes with m6A modifications that peaked at different stages in the polysome-occupied dataset and analyzed their biological functions (**Fig. 4F**). These m6A-modified genes were involved in key processes such as chromosome segregation, mitotic division, and RNA splicing. Together, these findings underlie the importance of m6A marks translationally active transcripts.

### Functional validation reveals 2-Cell-specific translation of eIF1ad3 and its variants are essential for embryo development

With this unpreceded genome-wide translational profiles of pre-implantation development, we have also functionally validated the identified key regulators specifically involved in the EGA. From the list of genes that are translationally activated during the 2-Cell stage (**Fig. 2F**), we identified a group of genes, including *Eif1ad2, Eif1ad3, Eif1ad4, Eif1ad6, Eif1ad7, Eif1ad8, Eif1ad16* and *Eif1ad19,* whose roles in embryogenesis remain unknown (**Fig. 5A**). These genes, in particular Eif1ad2, Eif1ad3, Eif1ad4, Eif1ad6, Eif1ad7 and Eif1ad8, have highly conserved sequences with only minimal nucleotide mismatches, but differ from well-studied translation initiation factors Eif1 and Eif1ad (F**ig. 5B****, Fig. S8A**). We therefore focused on targeting Eif1ad3 and its variants to investigate their function in mouse pre-implantation development. Due to the high sequence similarity among these genes, we were unable to specifically knock down a single gene to study the function of one specific gene.

**Figure 5.**
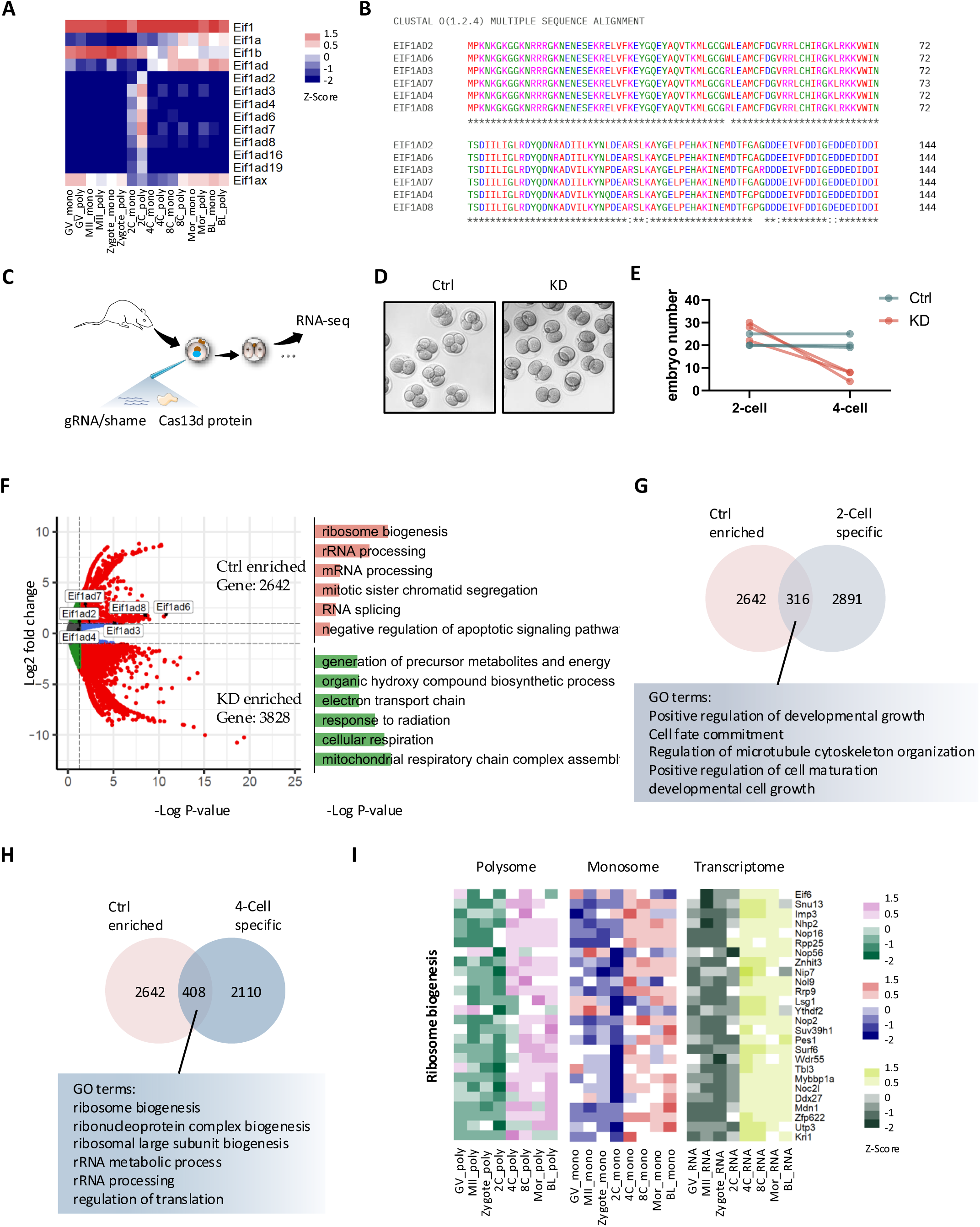
2-Cell-specific translation of eIF1ad3 and its variants are essential for embryo development. **A.** Heatmap showing the expression level of Eif1 family genes. The color spectrum, ranging from red through white to blue, indicates high to low levels of gene expression. **B.** Multiple sequence alignment (Omega) of Eif1ad amino acid sequence. **C.** Experimental scheme of injecting gRNA and RfxCas13d mRNA into zygote stage mouse embryos. **D.** A representative image of mouse embryos from control and knockdown groups. **E.** 4-Cell developmental rate of Eif1ad3 knockdown embryos compared to control group. **F.** Volcano plots showing the number of up- or down-regulated genes in control compared to Eif1ad knockdown at 2-Cell stage, the top GO terms of up- and down-regulated genes are shown separately. **G.** Venn diagram showing the number of genes and their top representative GO terms that are overlapped between control enriched and 2-Cell specific genes from polysome profile. **H.** Venn diagram showing the number of genes and their top representative GO terms that are overlapped between control enriched and 4-Cell specific genes from polysome profile. **I.** Heatmap in polysome profile, monosome profile, and transcriptome profile showing the expression level of genes regulating ribosome biogenesis.

By using the CRISPR/RfxCas13d system to knockdown (KD) Eif1ad3 variants (**Fig. 5C**), we found that almost all embryos in the KD group arrested at the 2-Cell stage (**Fig. 5D, E**). RNA-seq analysis of individual embryos (both KD and control) at the 2-Cell stage revealed that 2,642 genes were highly enriched in the control group. They were mainly related to ribosome biogenesis, RNA processing and cell division, which are essential for embryo activation during EGA (**Fig. 5F**). In contrast, 3,828 genes that are enriched in the KD group were associated with mitochondrial function, cellular respiration and metabolite formation, indicating apoptosis and embryo death (**Fig. 5F**). By comparing stage-specific translated genes from **Fig. 2D** (cluster 6-8) with the control-enriched genes, we identified 316 common genes that are involved in developmental growth, cell fate commitment, and cell maturation (**Fig. 5G**). During EGA transcription, new mRNAs are synthetized and at the same time translation is reprogrammed via ribogenesis ^10^. It is obvious that the mRNAs associated with this process are in translation at polysomes (**Fig. 3F**). In fact, the top represented GO terms including ribosome biogenesis and rRNA processing that enriched in control groups were found in the overlapped genes specific in 4-Cells instead of 2-Cells (**Fig. 5H, I**). Ribosome assembly, one of the major active biological processes from 4-Cells until blastocysts (**Table 1**), is impaired in the Eif1ad3 KD embryos, leading to the developmental arrest.

### Eif1ad3 regulates translation of functional genes during EGA

Eukaryotic initiation factors (eIFs) play a crucial role in initiation of translation ^38,39^. To investigate the interaction of Eif1ad3 with endogenous RNA in 2-Cell embryos, we injected *in vitro* transcribed Eif1ad3 mRNA with a His-tag (Eif1ad3-His IVT-mRNA, **Fig. 6A, B**) into zygotes due to the lack of Eif1ad3 specific antibody. Eif1ad3-His do not affect development up to the blastocyst stage (**Fig. 6C, D**). We then performed RIP-seq on 2-Cell stage embryos using a His-tag antibody and compared the results between the Eif1ad3-His group and a control IgG group. PCA results showed clear separation between groups, except for IgG, likely due to nonspecific interactions (**Fig. S8B**). After eliminating nonspecific interactions, we identified 3,708 genes directly or indirectly associated with Eif1ad3, indicating its broad involvement in initiating mRNA translation during EGA (**Fig. 6E**). These genes were primarily involved in processes such as Golgi vesicle transport, RNA processing, RNA modification, and protein maturation, all closely related to translation (**Fig. 6F**). Additionally, several ribosomal proteins RNAs were significantly bound to Eif1ad3, suggesting its role in ribosome assembly (**Fig. 6G**).

**Figure 6.**
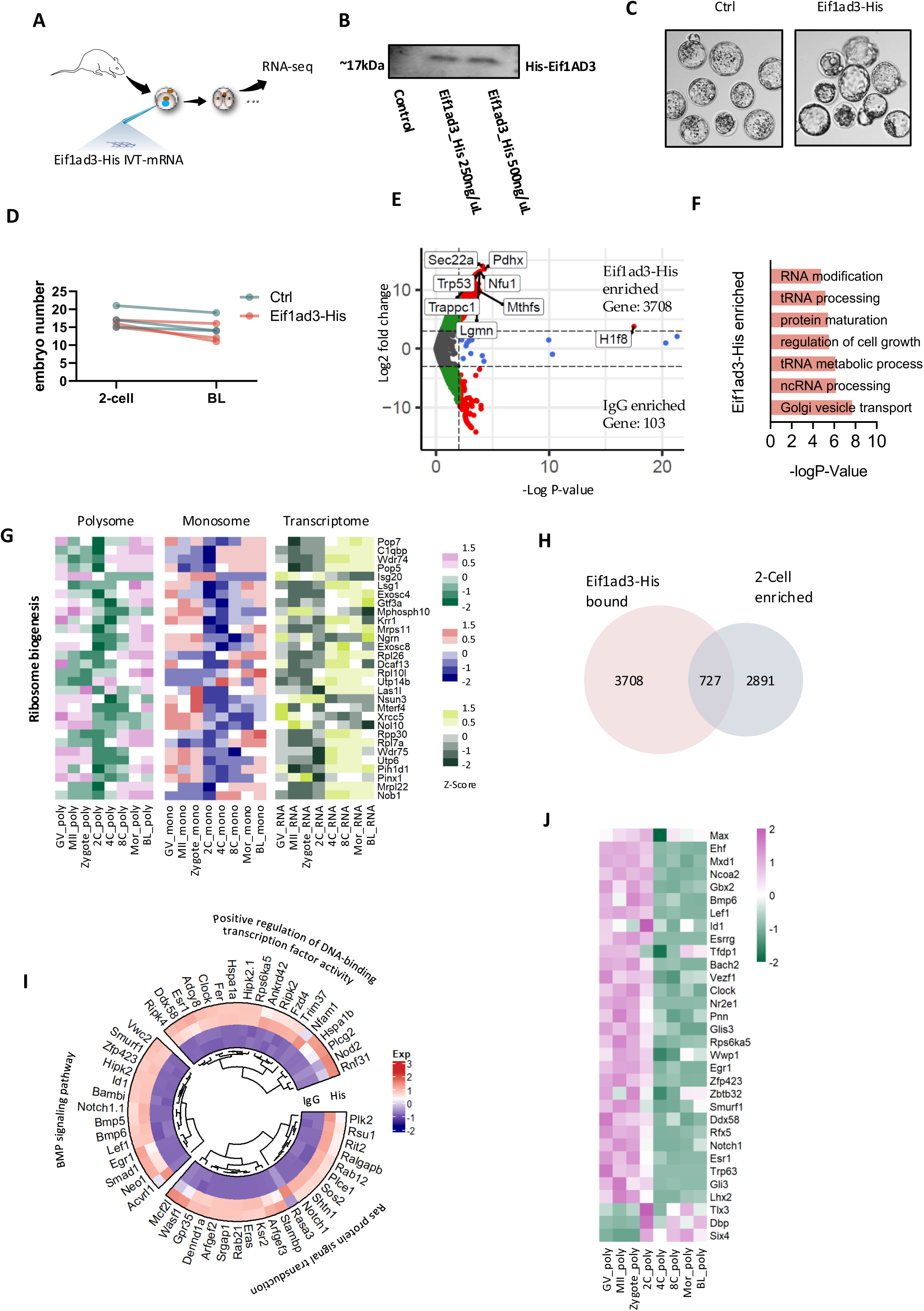
Eif1ad3 regulates translation of functional genes during EGA. **A.**Experimental scheme of injecting Eif1ad3-His mRNA into zygotes. **B.** Representative protein levels of His after injecting Eif1ad3-His mRNA at different concentrations into mouse zygotes. **C.** Representative image of mouse blastocysts from ctrl Control and Eif1ad3-His groups. **D.** Quantification of blastocyst rates of embryos with Eif1ad3-His compared to control. **E.** Volcano plots showing the number of up- or down-regulated genes in RIP-His compared to RIP-IgG group at 2-Cell stage. **F.** Main biological functions regulated by RIP-His group enriched genes. **G.** Heatmap in polysome profile, monosome profile, and transcriptome profile showing the expression level of genes regulating ribosome biogenesis. **H.** Venn diagram showing the number His-enriched genes from RIP-seq compared to 2-Cell peaked genes from polysome profile. **I.** Representative GO terms enriched from overlapped genes between His-enriched genes from RIP-seq and 2-Cell peaked genes from polysome profile, as well as the genes in each term. The color spectrum, ranging from red through white to blue, indicates high to low levels of gene expression. **J.** Heatmap showing the expression level of transcription factors within overlapped genes between His-enriched genes from RIP-seq and 2-Cell peaked genes from polysome profile. The color spectrum, ranging from pink through white to green, indicates high to low levels of gene expression.

We further narrowed down genes that are enriched in the 2-Cell stage from polysome profiles (**Fig. 2A**, Clusters 6-8) to 727 genes (**Fig. 6H**). These genes regulate DNA-binding transcription factor activity and affect BMP and Ras signaling pathways (**Fig. 6I**). Since transcription factors (TFs) are crucial for genome activation across mammalian species ^40,41^, we analyzed the TFs within these 727 genes (**Fig. 6J**). These TFs were highly involved in EGA, including BMP signaling, cell proliferation, and cell fate commitment (**Fig. S8C**).

In summary, Eif1ad3 is essential for EGA during mouse pre-implantation development by regulating the translation of mRNAs encoding factors that maintain developmental competence.

## DISCUSSION

The mRNA translation landscape and particularly the translational controls operating on specific mRNAs during rapid period of pre-implantation development is poorly understood. Although Ribo-seq has been used to characterize the translational landscapes of mouse oocytes and pre-implantation embryos ^10,11,42^. However, these analyses preclude exploration of the spatial landscape of the dynamic mRNA translation process due to the lack of single-fraction resolution of ribosome-occupied RNAs. Here, by utilizing a low-input, high-resolution, optimized SSP-profiling ^12^ polysome profile integrated with RNA-seq, we were able to overcome this critical limitation. Instead of exploring ribosome-occupied RNA as a single profile, we analyzed ten individual fractionation profiles based on the number of bounding ribosomes. It not only significantly increased the depth of the sequencing, but also enabled us to track the dynamics of translational fate for mRNAs. The spatial map of mRNA translational fate represents another level of translational regulation and is critical for understanding the mechanisms and strategies that oocyte and embryo employ to rapidly respond to the demands arising from successive but distinct stages during pre-implantation development. Furthermore, by comparing polysomal and monosomal-occupied mRNA profiles, we have identified critical translationally activated genes that contribute to oocyte maturation, oocyte-embryo transitions, EGA, and lineage specification.

Our study was able to capture diverse, although somewhat empirical, modes of translational selectivity for transcripts. Our results suggest that the three major mRNA fates presented in oocytes and early embryos are indicative of mechanisms employed by embryos to effectively reduce energy consumption and the time required for transcript re-synthesis, likely reflecting the requirements of translational dynamics during pre-implantation embryo development. Maternal mRNAs are known to be important for pre-implantation development ^40^. However, it remains unclear whether translation of specific maternally stored mRNAs is essential for EGA or later stages. Here, by analyzing the genes from Mode 1, we identified a group of maternal RNAs that bind to the monosome at the zygote stage but exclusively to the polysome at the 2-Cell stage (**Fig. 2E top panel**), undoubtedly indicating their specific functions during EGA. Furthermore, this database enabled us to reveal the fate of each maternal mRNA across the different pre-implantation stages: It is either degraded without ribosome binding, stored by monosome binding or translated upon polysome occupancy. A comparative analysis of datasets from oocytes and pre-implantation embryos occupied with ribosomes and monosomes showed dynamic changes in the stage-specific activated translatome landscape. Interestingly, only a small fraction of polysome-enriched RNAs were found to maintain this status across all stages, especially after the zygote stage. Most of the polysome-enriched RNAs are activated in a stage-specific manner by groups with significant and non-significant levels of monosome enrichment, reflecting precise regulation for the development of the pre-implantation embryo.

The length of poly(A) tails, which varies widely across different mRNAs, has a significant impact on both their stability and translational activity ^31,43^, therefore plays a crucial role in coordinating the translation during maternal-zygotic transition. Additionally, polyadenylation, occurring at different stages, helps fine-tune gene expression by selectively stabilizing certain transcripts while allowing others to degrade ^29,30^. Our study highlighted a notable delayed positive correlation between poly(A) tail length at the 2-Cell stage and polysome association at the 4-/8-Cell stages. This suggests that while poly(A) tails are elongated at the 2-Cell stage, their functional impact on translation becomes more evident in subsequent stages of development. Furthermore, this delay could indicate a temporal control mechanism, where poly(A) tail lengthening prepares transcripts for translation only after key developmental checkpoints are passed, ensuring precise expression in the highly dynamic activation of specific mRNAs. This observation aligns with other studies that highlight poly(A) tail modifications as a form of translational control that governs early embryonic development and establishes the foundation for cell lineage decisions ^44,45^.

We have identified a key eukaryotic initiation factor, Eif1ad3, that is actively translated exclusively at the 2-Cell embryos and essential for EGA during mouse embryonic development. eIFs are fundamental to the translation initiation process and act as primary regulators of gene expression patterns during development. While it has been found a number of genes in the eIF family play essential roles in the mouse pre-implantation development including eIF1A^46^, eIF3^47^, eIF4E^48^, and eIF4E1b ^49^, we extended this list by providing the essential role of other eIF factors, Eif1ad3 and its variants, in the mammalian pre-implantation development. Interestingly, Eif1ad3 exhibits a translation pattern distinct from Eif1a and Eif1ad and is specifically translated at the 2-Cell stage. Deficiency of Eif1ad3 leads to embryonic arrest at the 2-Cell stage. Surprisingly, the large number of genes affected by Eif1ad3 knockdown regulates ribosome biogenesis and associated rRNA synthesis. These genes are only marginally transcribed but are extensively translated from the 4-Cell stage, which is consistent with the transition from the maternal to the embryonic stage. Consequently, the 2-Cell arrest of embryos could be due to a lack of ribosome formation required for translation reprograming post-EGA, warranting further investigation.

In summary, our study has revealed a previously unappreciated level of mRNA translational fate associated with mouse oocyte maturation and embryo development. We have shown that different maternal mRNAs undergo distinct fates and modifications at different preimplantation stages by being either degraded, stored or actively translated, which are crucial for embryos to efficiently manage energy and prepare for successive developmental steps. The essential roles of 2-Cell specific translated genes, *Eif1ad3* and its variants in mouse early development was particularly striking observations, further demonstrating significant value of this unpreceded resource on prioritizing genes for functional characterizations.

## MATERIALS AND METHODS

### Animal care and use

All animal experiments were performed in accordance to guidelines and protocols approved by the University of Florida Institutional Animal Care and Use Committee (IACUC) under protocol number IACUC202300000558, and the Laboratory of Biochemistry and Molecular Biology of Germ Cells at the Institute of Animal Physiology and Genetics in Czech Republic under Act No. 246 / 1992 on the protection of animals against cruelty, issued by the Ministry of Agriculture experimental project #67756 / 2020MZE-18134.

#### Mice

ICR females (produced in house, IAPG, Czech Republic) and CF-1 females (Envigo, Indianapolis, IN, USA) used for oocytes and embryo collection for the SSP profiling and knock down experiments. B6SJLF1/J males used for breeding were obtained from Jackson Laboratory.

#### Antibodies and reagents

A list of all primary antibodies, reagents, as well as Oligonucleotides used in this study can be found in Supplementary Table S2.

### Oocyte and embryo isolation and cultivation

Oocytes were acquired from ICR mice of a minimum of 6 weeks old. The females were stimulated 46 h prior to oocyte isolation using 5 IU of pregnant mare serum gonadotropin (PMSG; Folligon; Merck Animal Health) per mouse. Fully grown GV oocytes were isolated into transfer medium (TM) supplemented with 100 μM 3-isobutyl-1-methylxanthine (IBMX, Sigma Aldrich) to prevent spontaneous meiotic resumption. Selected oocytes were denuded and cultivated in M16 medium (Merck Millipore) without IBMX at 37 C, 5% CO 2 for 0 (GV) or 12 h (MII). For embryo collection, the PMSG stimulated mice were injected with 5 IU hCG before being mated overnight with males of the same strain. After 16 h, zygotes were recovered from the excised oviducts and cultured in KSOM medium (Merck Millipore) until blastocyst stage. Interphase pronuclei zygotes were collected at the time point of isolation and later stages 4-Cell, 8-Cell, morula and blastocyst stages were collected subsequent days.

### Isolation of ribosome-bound mRNA

Approximately 200 oocytes (GV or MII oocyte) or embryos at different developmental stages (zygote, 2-, 4, 8-Cell, morula, and blastocyst) were combined with lysis buffer containing 10 mM HEPES (pH 7.5), 5 mM KCL, 5 mM MgCL2, 2 mM DTT, 1% Triton X-100, 100 μg/ml cycloheximide, complete EDTA-free protease inhibitor (Roche), and 40 U/ml RNase inhibitor. Oocytes and embryos were disrupted by zirconium silica beads (Sigma) in the mixer mill apparatus MM301 (shake frequency 30, total time 45s, Retsch). Lysates were cleaned by centrifugation in 10,000 x g for 5 min at 4°C and the supernatants were loaded into 10-40 % linear sucrose gradients containing 10 mM HEPES (pH7.5), 100 mM KCl, 5 mM MgCl2, 2mM DTT, 100 μg/ml cycloheximide, complete EDTA-free protease inhibitor, and 5 U/ml RNase inhibitor. Ultracentrifugation was carried out with a SW55Ti rotor and Optima L-90 Ultracentrifuge (Beckman Coulter). Ribosome profiles were recorded by ISCO UV absorbance reader. Ten equal fractions were then recovered and subjected to RNA isolation by Trizol reagent (Sigma).

### Library preparation and RNA sequencing (RNA-seq)

The RNA-seq libraries were generated from individual fractions by using the Smart-seq2 v4 kit with minor modification from manufacturer’s instructions. Briefly, individual cells were lysed, and mRNA was captured and amplified with the Smart-seq2 v4 kit (Clontech). After AMPure XP beads purification, amplified RNAs were quality checked by using Agilent High Sensitivity D5000 kit (Agilent Technologies). High-quality amplified RNAs were subject to library preparation (Nextera XT DNA Library Preparation Kit; Illumina) and multiplexed by Nextera XT Indexes (Illumina). The concentration of sequencing libraries was determined by using Qubit dsDNA HS Assay Kit (Life Technologies) and KAPA Library Quantification Kits (KAPA Biosystems). The size of sequencing libraries was determined by means of High Sensitivity D5000 Assay in at Tapestation 4200 system (Agilent). Pooled indexed libraries were then sequenced on the Illumina HiSeq X platform with 150-bp paired-end reads.

A pool of 20 oocytes or preimplantation embryos (n=4) selected from the same batch in each developmental stage used for ribosome profiling were used to profile transcriptomes by RNA-seq following Smart-seq2 protocol as above described.

### RNA-seq data analysis

Multiplexed sequencing reads that passed filters were trimmed to remove low-quality reads and adaptors by TrimGalore (v.0.6.7) (-q 25 --length 20 --max_n 3 --stringency 3). The quality of reads after filtering was assessed by fastQC, followed by alignment to the mouse genome (GRCm39) by HISAT2 (v.2.1.0) with default parameters. The output SAM files were converted to BAM files and sorted using SAMtools (v.1.17). Read counts of all samples were quantified by using FeatureCount (v.2.0.1) with mouse genome as a reference and were adjusted to provide CPM (Counts per million mapped reads). Principal component analysis (PCA) and cluster analysis were performed by using R (a free software environment for statistical computing and graphics). Differentially expressed genes (DEGs) was identified by using DESeq2 (v.4.2.1) in R. The genes were considered differentially expressed if they provided a false discovery rate of <0.05 and fold change >2. Clustering of time series gene expression data was performed by using Mfuzz (v.3.19) in R. ClusterProfiler (v.4.6.1) was used to reveal the Gene Ontology (GO) and KEGG pathways in R.

### Inhibitor treatment

Cells were treated with 10uM of W-7 hydrochloride at 2Cell stage, and untreated cells were considered as control group (Initial n=80* for each group, considered as 100% in the graph). Embryos kept in the presence of the inhibitor for the rest of the experiment, and the development status checked for each stage.

### Knockdown in mouse embryos

The microinjection platform was equipped with a microinjector (FemtoJet 4i, Eppendorf), an inverted microscope (Leica, DMi8), and micromanipulators (Narishige). Microinjections were performed between 20 and 24 hours after hCG administration. For the Eif1ad3 knockdown experiments, 1-cell embryos were collected from CF1 (Envigo) mice and microinjected with either control (RfxCas13d-IVT mRNA, 500 ng/µL) or Eif1ad knockdown (gRNA, 2 µM + RfxCas13d-IVT mRNA, 500 ng/µL). Embryos from both control and knockdown groups were cultured to the blastocyst stage to observe development. For RNA-Seq, 2-cell stage embryos (from both control and knockdown groups) were collected.

### In vitro transcription of Eif1ad3

To generate Eif1ad3-6xHis mRNA, total RNA from 2-Cell embryos was extracted from a pool of mouse embryos, followed by first strand cDNA synthesis using SuperScript™ IV VILO™ Master Mix (Thermo Fisher Scientific, Waltham, MA). Primers were designed based on the current genome annotation (GRCm39; National Center for Biotechnology Information) of Eif1ad3 (Table S2). PCR was conducted using Q5 Hot Start High-Fidelity 2X Master Mix (New England Biolabs, Ipswich, MA) with an initial denaturation step at 98°C for 30 seconds followed by 30 cycles at 98°C for 10 seconds, annealing at 58°C for 30 seconds and extension at 72 °C for 30 seconds and a final extension at 72 °C for 2 minutes. The purified PCR products were served as DNA template for in vitro transcription using HiScribe® T7 ARCA mRNA Kit with tailing following manufacture’s instruction. The yield and integrity of resulting mRNA were assessed using Qubit 4 (Thermo Fisher Scientific, Waltham, MA) and Tapestation 4150 (Agilent Technologies, Santa Clara, CA). To overexpress Eif1ad3, in vitro transcribed mRNAs (IVT-mRNAs) were microinjected into zygotes at final concentration of 200 ng/uL.

### RNA Immunoprecipitation Sequencing (RIP-seq)

RIP experiments were conducted using the EZ-Magna RIP Kit (Millipore Corporation, Billerica, MA). Briefly, 2-Cell stage embryos (over expressed 200 cells and control 200 cell per replicate) were harvested, lysed and then incubated with RIP buffer containing magnetic beads conjugated with anti-His tag antibody (Millipore) or corresponding negative control IgG (Millipore). After the antibody was recovered using protein A/G beads, the purified RNA were used to perform RNA-seq.

### Immunoblotting

An exact number of cells (15–30 oocytes) were washed in PVA / PBS and frozen to −80 ZC. Prepared samples were lysed in NuPAGE LDS Sample Buffer (NP0007, Thermo Fisher Scientific) and NuPAGE Sample Reducing Agent (NP0004, Thermo Fisher Scientific) and heated at 100 ZC for 5 min. Proteins were separated on precast gradient 4–12% SDS–PAGE gel (Thermo Fisher Scientific) and blotted to Immobilon P membrane (Millipore) in a semidry blotting system (Biometra GmbH) at 5 mA cm 2 for 25 min. Membranes were then blocked in 5% skimmed milk dissolved in 0.05% Tween-Tris buffer saline (TTBS), pH 7.4 for 1 h. Membranes were incubated overnight at 4 C with relevant primary antibodies (Table S2) diluted in 1% milk / TTBS. Appropriate Peroxidase conjugated secondary antibodies were used (711-035-152 Anti-Rabbit Donkey, or 715-035-151 Anti-Mouse Donkey, both Jackson Immuno research) at a 1:7500 dilution in 1% milk / TTBS for 1 h at room temperature. ECL (Amersham) was used for visualization of immunodetected proteins on Azure Biosystems.

### Quantification and statistical analysis

Statistical differences between pairs of datasets were analyzed by two-tailed unpaired t-tests. Values of p < 0.05 were considered statistically significant. Where applicable, all quantitative data are presented as the mean ± SEM. Repeated number was indicated as ‘‘n’’ in figure legends.

## Data availability

The raw FASTQ files and normalized read accounts per gene are available at Gene Expression Omnibus (GEO) (https://www.ncbi.nlm.nih.gov/geo/) under the accession number GSE263902 and GSE279822. The previously published datasets including proteomics (PXD003315), polyA (GSE228001) and m6A modification (GSE192440) were downloaded from ProteomeXchange Consortium and SRA database and re-analysed in this study. The mouse reference genome GRCm39 (Release 113) was downloaded from the Ensembl (https://ftp.ensembl.org/pub/release-113/fasta/mus_musculus/).

## Acknowledgements

This work was supported by the NIH Eunice Kennedy Shriver National Institute of Child Health and Human Development (R01HD102533) and USDA National Institute of Food and Agriculture (2019-67016-29863, W4171, Capacity fund hatch project 7004459), Institutional Research Concept RVO67985904 and The Czech Science Foundation (22-27301S), and MSMT (EXCELLENCECZ.02.1.01/0.0/0.0/15_003/0000460 OP RDE).

## Author contributions

Z.J., and A.S. designed and supervised the research. H.M., and R.I. conducted research, analyzed the data, and interpreted and assembled the experimental results. K.K., and M.D. performed the western blot validation experiments. H.M., R.I., A.S., and Z.J. wrote the manuscripts.

## Supplementary Figures

**Figure S1. Principal Component Analysis (PCA) of sequencing data.** A. PCA of 4 dimensions (PC1-4) of 10 fractions of mouse oocytes and early embryos. B. Principal Component Analysis (PCA) of free-, monosome- and polysome-bound mRNA profiles in 10 fractions of mouse oocytes and early embryos.

**Figure S2. Flow of genes across modes during mouse oocyte and preimplantation development. A-D.** Expression dynamic of candidate mRNAs from polysome profile, monosome profile, and transcriptome profile.

**Figure S3. Comparison of ribosome-bound and ribosome-unbound mRNA in oocytes and early embryos. A.** Expression dynamic of selected genes from polysome profile, monosome profile, and free RNA profile. **B.** Heatmap in both polysome profile and monosome profile showing the expression level of genes from cluster7 in Figure 2A. The color spectrum, ranging from red through white to blue or from green through green to pink, indicates high to low levels of gene expression. **C.** Dot plot comparison showing top GO terms enriched from stage-specific genes from monosome and polysome datasets. Color from grey to red represents the p-value from high to low, size from small to large represents gene number from low to high. **D.** Dot plot comparison showing top GO terms enriched from stage-specific genes from free RNA datasets. The color from grey to red represents the p-value from high to low, dots from small to large represents gene number.

**Figure S4. Trajectory of ribosome bound mRNA during mouse oocyte and preimplantation development. A.** Representative overlapped genes peaking from GV till 2-Cell stage from monosome-bound and polysome-bound profiles, as well as the biological functions being regulated correspondingly. **B.** Representative overlapped genes peaking from 4-Cell till blastocyst stage from monosome-bound and polysome-bound profiles, as well as the biological functions being regulated correspondingly.

**Figure S5.** Expression dynamic of candidate mRNAs from polysome profile, monosome profile, and free RNA profile.

**Figure S6. Genes essential for critical fate transitions during mouse oocyte and early embryo development. A.** Heatmap of critical genes translationally activated in GV to MII transition. **B.** Heatmap of critical genes translationally activated in zygote to 2-Cell transition. **C.** Heatmap of critical genes translationally activated in 8-Cell to blastocyst transition. The color spectrum, ranging from red through white to blue or from green through green to pink, indicates high to low levels of gene expression.

**Figure S7.** Correlation between polysome-occupied RNA level (4-/8-Cell) and protein level (morula).

**Figure S8. Injecting Eif1ad3-His into mouse zygote does not affect embryo development. A.** Schematic representation of Eif1a, Eif1ad, and Eif1ad3 amino acid sequence. **B.** PCA of samples of Eif1ad3-His, IgG, negative control, and total RNA groups from RIP-seq. **C.** Representative GO terms enriched from the transcription factors. Color from pink to green represents the p-value from low to high, size from small to large represents gene number from low to high.

