## Supplementary material for "Spatiotemporal dynamics and selectivity of mRNA translation during mouse pre-implantation development": Figure S8

Fig. S8

A

```
CLUSTAL O(1.2.4) multiple sequence alignment


Eif1_NP_035638.1            -------------MSAIQNLHSFDPFADASKGDDLLPAGTEDYIHIRIQQRNGRKTLTTV        47
Eif1ad3_NP_001116134.1      MPKNKGKGGKNRRRGKNENESEKRELVFKEYG--------QEYAQVTKMLGCGRLEAMCF        52
Eif1ad_NP_081512.1          ----------------MSQASKRKHVVQEVLGEHMVPSDHQQIVKVLRTPGNNLHEVETA        44
                                             .:  .   ..    *        ::  ::      .

Eif1_NP_035638.1            QGIA------DDYDKKKLVKAFKKKFACNGTVIEHPEYGEVIQLQGDQRKNICQFLIEIG        101
Eif1ad3_NP_001116134.1      DGVRRLCHIRGKLRKKVWINTS--DI----ILIGLRDYQ-D---------NKADVILKYN        96
Eif1ad_NP_081512.1          QGQRFLVSMPSKYRKNIWIKRG--DF----LIVDPIEEGEK---------VKAEISFVLC        89
                            :*         ..  *:  ::     .:      ::   :               .:. :

Eif1_NP_035638.1            LAKDDQLKVHGF------------------------------------------------        113
Eif1ad3_NP_001116134.1      PDEARSLKAYGELPEHAKI-----NEMDTFGAR-DDDEIVFDDIGEDDEDIDDI------        144
Eif1ad_NP_081512.1          KNHVRSLQKEGHWPEAFSEVAEKQNNMNRESQPELPAEPQLSGEGSSSEDDSDLFVNTNH        149
                              .  .*:  *

Eif1_NP_035638.1            ---------------------       113
Eif1ad3_NP_001116134.1      ---------------------       144
Eif1ad_NP_081512.1          RQYHESEEESEEDEEEEEEAA       170
```

B

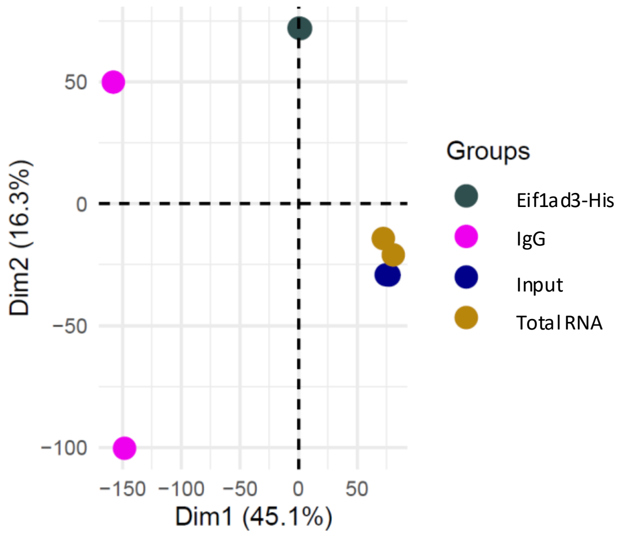


C

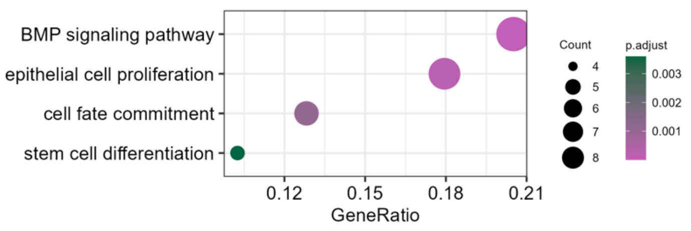
